# Amyloid fibril polymorphism in the heart and liver of an ATTR amyloidosis patient with polyneuropathy attributed to the ATTRv-V122Δ variant

**DOI:** 10.1101/2024.05.09.593396

**Authors:** Yasmin Ahmed, Binh An Nguyen, Candace Kelly, Shumaila Afrin, Virender Singh, Bret M. Evers, John M. Shelton, Christian Lopez Escobar, Preeti Singh, Rose Pedretti, Lanie Wang, Parker Bassett, Maria del Carmen Fernandez-Ramirez, Maja Pekala, Andrew Lemoff, Barbara Kluve-Beckerman, Lorena Saelices

## Abstract

ATTR amyloidosis is a phenotypically heterogeneous disease characterized by the pathological deposition of transthyretin in the form of amyloid fibrils into various organs. ATTR amyloidosis may result from mutations in variant (ATTRv) amyloidosis, or aging in wild-type (ATTRwt) amyloidosis. ATTRwt generally manifests as cardiomyopathy, whereas ATTRv may present as polyneuropathy, cardiomyopathy, or mixed, in combination with many other symptoms deriving from multisystem organ involvement. Over 220 different mutational variants of transthyretin have been identified, many of them being linked to specific disease symptoms. Yet, the role of these mutations in explaining differential disease manifestations remains unclear. Using cryo-electron microscopy, here we structurally characterized fibrils from the heart and the liver of an ATTRv patient carrying the V122Δ mutation, which is predominantly associated with polyneuropathy. Our results show that these fibrils are polymorphic, presenting as both single and double filaments. Our study alludes to a structural connection contributing to phenotypic variation in ATTR amyloidosis, as polymorphism in ATTR fibrils may manifest in patients with predominantly polyneuropathic phenotypes.

**Significance:** ATTR amyloidosis is a systemic, clinically diverse disease that results in organ failure due to the accumulation of transthyretin amyloid fibrils. ATTR patients present with varied symptoms, yet the root of this phenotypic heterogeneity remains unclear. Previous studies suggest an association between phenotype and fibril structure polymorphism. Here we describe the cryo-electron microscopy structure of variant transthyretin amyloid fibrils associated with a predominantly polyneuropathy phenotype. We have found polymorphism within these fibrils, a phenomenon we have thus far only observed in polyneuropathic associated transthyretin mutations. Our results signify an association between fibril structure and phenotype in ATTR amyloidosis.

## Introduction

Transthyretin amyloidosis (ATTR) amyloidosis is a fatal, phenotypically diverse disease characterized by the deposition of amyloidogenic transthyretin (ATTR) fibrils in various organs. Transthyretin is mainly synthesized by the liver, although other tissues secrete it in lower amounts, including the retinal pigment epithelium and the choroid plexus^1^. ATTR deposition may stem from mutations in the *TTR* gene in variant ATTR (ATTRv) amyloidosis, or by unknown aging-related factors in wild-type ATTR (ATTRwt) amyloidosis^2^. While ATTRwt amyloidosis is clinically more predictable, ATTRv amyloidosis often presents with complex and variable clinical manifestations including polyneuropathy, cardiomyopathy, or a mixed phenotype, in addition to many other symptoms stemming from secondary organ involvement including carpal tunnel syndrome, gastrointestinal dysfunction, and ocular issues^3^. More than 220 mutational variants of transthyretin have been documented, with many showing associations for specific disease symptoms (∼93%)^4^. The influence of each mutational variant on the clinical phenotype is not yet fully understood, and some studies suggest that it could be related to their effects on fibril formation, structure, or both^5,6^.

Recent cryo-electron microscopy (cryo-EM) studies have contributed important structural insights into the amyloid assemblies of various diseases^7–9^. Notably, studies on tau, α-synuclein, amyloid-β, and TDP-43, have all revealed disease-specific fibril polymorphism, in which a particular fibrillar fold is linked to a specific disease^10–14^. Exemplifying this phenomenon, Alzheimer’s disease patients exhibit the same two types of tau fibrils: paired helical filaments and straight filaments. And yet, these structures differ from the ones adopted by tau in chronic traumatic encephalopathy or other tauopathies^15^. Another example is found in amyloid-β filaments from Alzheimer’s disease patients. Amyloid-β filaments can adopt several different amyloid folds within the same individual—nonetheless, the structures remain specific to the disease^14^.

In ATTR amyloidosis, cryo-EM investigations have revealed a substantial structural variability whose influence on phenotype remains to be established. Early cryo-EM studies of ATTR fibrils revealed a unique structural conformation, made of two fragments of transthyretin and forming a polar channel that runs through the fibril^16^. We and others have found that the amyloid fibrils from multiple ATTRwt amyloidosis patients are structurally homogenous, which is consistent with their more uniform disease presentation as cardiomyopathy^16,17^. In contrast, our recent study on ATTR fibrils from polyneuropathic ATTRv-I84S patients reveals an unprecedented structural polymorphism^18^, as each of the three patients had two populations of fibrils: one that was shared by all three patients, similar to the one observed in the early studies; and one that was specific to each individual, with local structural variations in the polar channel. The nature and location of the structural variations suggest that the mutation I84S may drive fibril polymorphism in these patients. Additionally, previous studies have disclosed that the amyloid fibrils deposited in the heart^16^ and the vitreous humor of the eye^19^ of ATTRv-V30M patients can adopt distinct structures that differ at two levels: the conformation of the fibril has local variations in the polar channel and the fibrils have different numbers of protofilaments. These studies suggest that ATTR fibril polymorphism may also stem from the organ of deposition and/or the source of transthyretin. Additional structural investigations are necessary to ascertain the potential role of ATTR fibril polymorphism in clinicopathological heterogeneity, should there be any, and to elucidate the underlying causes of structural variability.

The primary aim of this study, therefore, is to expand fibril structural data for polyneuropathic ATTR amyloidosis, complementing existing data from cardiomyopathic cases. Understanding how fibril structural morphologies relate to clinical phenotypes is essential for improving the ATTR fibril targeted diagnostic tools and therapies. We used cryo-EM to determine the structure of fibrils extracted from the heart and the liver of an ATTRv-V122Δ patient^20^. This particular V122Δ mutation is the only documented deletion mutation in the *TTR* gene; and manifests predominantly as polyneuropathy, along with cardiomyopathy and carpal tunnel syndrome. Characterizing the fibrils structures of this mutation can provide an opportunity to investigate the potential structure – phenotype relationship in ATTR amyloidosis. Here, we show that ATTRv-V122Δ fibrils isolated from the heart and the liver tissues of a single patient with polyneuropathy are polymorphic, as they can ensemble into bundles made of single and double protofilaments. This study underlines the complexity of the structural landscape in ATTR amyloidosis and calls for further structural studies.

## Results

### Characterization of ATTR fibrils

Freshly frozen tissues were obtained from the explanted heart and liver of an Ecuadorian male heterozygous carrier of the ATTRv-V122Δ variant who became affected in his early 60s, and presented with both peripheral neuropathy and autonomic neuropathy^20^. Sections of the heart showed abundant deposits of amyloid within the interstitium of the myocardium, which was confirmed by Congo Red and Thioflavin-S staining (Fig. 1a-c). Examination of the epicardium revealed amyloid deposits within nerves, vessel walls, and adipose tissue (Fig. 1a-c). We isolated the ATTR fibrils from the heart and liver in accordance with a previously established mild water-based extraction procedure^16^. Mass spectrometry analysis confirmed transthyretin protein identity, ATTRv-V122Δ genotype, and the presence of both the mutant and wild-type transthyretin protein, as expected from this patient’s zygosity (Supplementary Fig. 1). We typed the sample by western blotting using an in-house antibody that specifically targets the C-terminal fragments of transthyretin (Supplementary Fig. 2a). These fragments are characteristic of type A ATTR fibrils, which are made of both full-length and C-terminal fragmented transthyretin, and distinguishes them from type B ATTR fibrils which are composed of solely full-length transthyretin^21^ (Supplementary Fig. 2a). The typing confirmed that the *ex-vivo* fibril extract from the ATTRv-V122Δ patient contained type A ATTR fibrils. We validated the amyloidogenic properties of the cardiac sample by examining its capacity to seed recombinant transthyretin aggregation *in vitro* (Supplementary Fig. 2b). We did not perform the seeding assay for the liver sample as the extraction purity was the major concern (Supplementary Fig. 2d). We verified the extraction of ATTR fibrils and their suitability for further cryo-EM analysis by negative staining transmission electron microscopy (Supplementary Fig. 2c-d).

**Figure 1:**
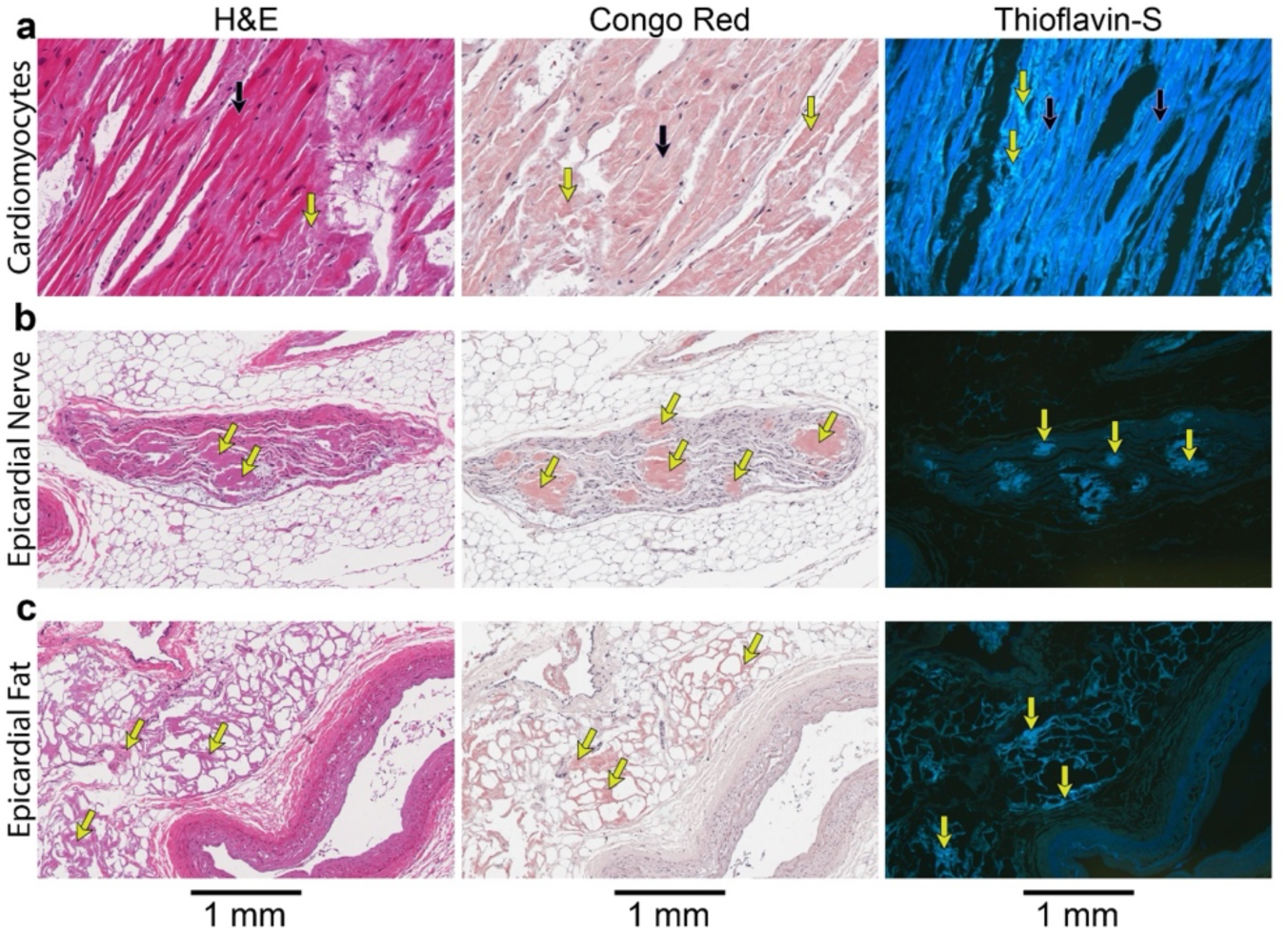
Histological analysis of ATTRv-V122Δ cardiac tissue. Hematoxylin and eosin (H&E), Congo Red, and Thioflavin-S staining (left to right) of different anatomical features of cardiac tissue, including cardiomyocytes (a), epicardial nerve (b), and epicardial fat. The staining depicts the localization and abundance of transthyretin amyloid deposits. Cardiomyocytes are denoted by black arrows, and the yellow arrows signify amyloid deposits. Scale bars, 1 mm.

### Structural polymorphism of ATTRv-V122Δ fibrils

We carried out cryo-EM analysis to determine the structure of ATTRv-V122Δ fibrils from both the heart and liver. For each sample, we optimized the fibril concentration and vitrification conditions to ensure optimal fibril distribution for cryo-EM data collection (Supplementary Fig. 2e-f). Fibrils were auto-picked using Topaz^22^. Two-dimensional (2D) classification of the collected images yielded multiple distinct morphologies. In the heart sample, we identified a minor population (4.34%) of straight fibrils lacking clear twists, as found in previous studies of ATTR fibrils, and thus they were unsuitable for helical reconstruction. The sample also included a major population (∼78.72%) of twisted single fibrils, similar in morphology to those found in other ATTR fibril extracts^16–19,23^ (Supplementary Table 1). To our surprise, we also found a smaller population (∼17%) of twisted double fibrils consisting of two intertwined protofilaments, which have not been observed in other cardiac ATTR fibrils to date (Fig. 2a, b). In the liver sample, we observed a similar distribution of morphologies: ∼6% straight fibrils, 81.7% twisted single fibrils, and 12.3% twisted double fibrils (Supplementary Fig. 3). The single and the double fibrils had distinct fibril widths as shown in Supplementary Fig. 4. We stitched the representative 2D class averages of single and double fibrils to estimate their respective crossover distances (Fig. 2b). We used these initial helical parameters to obtain three-dimensional (3D) classes. The density map of the single fibrils exhibited structural consistency with previously published ATTR fibrils (Fig. 2c)^16,17,23,24^. Using the 2D classification of the double fibrils, we built one initial model, low-pass filtered to 20 Å, and used that as the reference model for 3D classification (Fig 2b). The reconstruction of double fibrils revealed two distinct classes: one class with an apparent C2 symmetry (herein referred to as C-to-C double fibrils) and a second class with apparent C1 symmetry (herein referred to as N-to-C double fibrils). In both classes, each individual protofilament seemingly adopts the same structural conformation as in the single fibrils, but they differ in the inter-protofilament interface (Fig. 2c).

**Figure 2:**
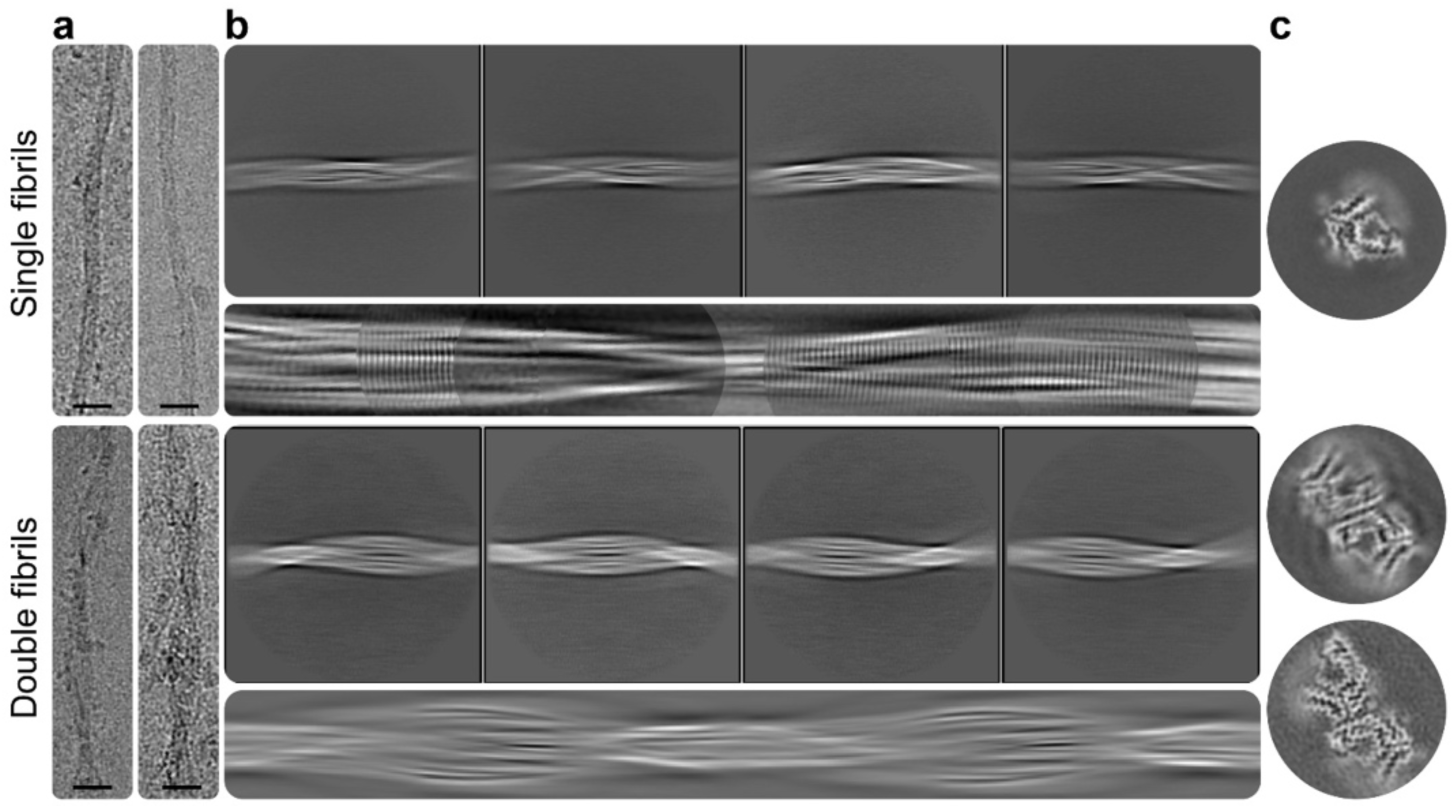
Cryo-EM data collection and processing of cardiac fibrils from an ATTRv-V122Δ patient. **a.** Representative cryo-EM micrographs of single (top) and double (bottom) fibrils. Scale bar, 20 nm. **b.** Representative 2D class averages of single (top) and double (bottom) fibrils extracted at particle box size of 768 pixel. Under the representative individual 2D classes panels, we include the stitching of all 2D classes of single fibril (top) and the initial model of double fibril generated in RELION (bottom). **c.** 3D class averages of single (top) and double (bottom) fibrils.

### Structure of the single ATTRv-V122Δ fibril closely resembles that of ATTRwt fibrils

We processed the 3D classes with single fibrils and generated density maps at resolution of 3.6 Å and 2.9 Å from the heart and liver extracts, respectively (Supplementary Fig. 5). Both the heart and liver fibril structures have similar crossover distances of 692 Å and 723 Å, a twist angle of - 1.27° and -1.19 (fibril was assumed to have left-handed twist), and a rise of 4.9 Å and 4.78 Å, respectively (Fig. 3a, b, Supplementary Fig. 6 & 7). The observed difference of the optimal rise was most likely due to calibration variations of the microscopes, because we collected data for the heart and the liver at two separate cryo-EM facilities. The reconstructed density map resembles the structure of cardiac ATTRwt^17^ and several ATTRv-V1fibrils previously solved. In detail, both models of ATTRv-V122Δ fibrils comprise two fragments: a N-terminal fragment spanning from Pro 11 to Lys 35 and a C-terminal fragment spanning from Leu 58 to Thr 122. Each layer of the fibril is stabilized by steric zippers throughout the entire structure, a hydrophobic pocket within the N-terminal fragment (17-LDAVRGSPAI-26), and an aromatic pocket within the C-terminal fragment consisting of Tyr78, Trp79, Phe87 and Phe 95. The residues Leu 58 to Ile 84 form a polar channel with no identifiable elements seen at the current resolution. The fibril structure is stabilized by backbone hydrogen bonds, intra-chain hydrogen bonds, and π-π stacking of aromatics residues. Structural comparison of the Cα with published ATTR structures (ATTRwt, ATTRv-I84S, ATTRv-V30M, ATTRv-V20I, ATTRv-G47E, and ATTRv-V122I) from previous studies and ATTRv-V122Δ demonstrates high similarity with a root mean square deviation (r.m.s.d) value of 0.65 Å (Fig. 3c).

**Figure 3:**
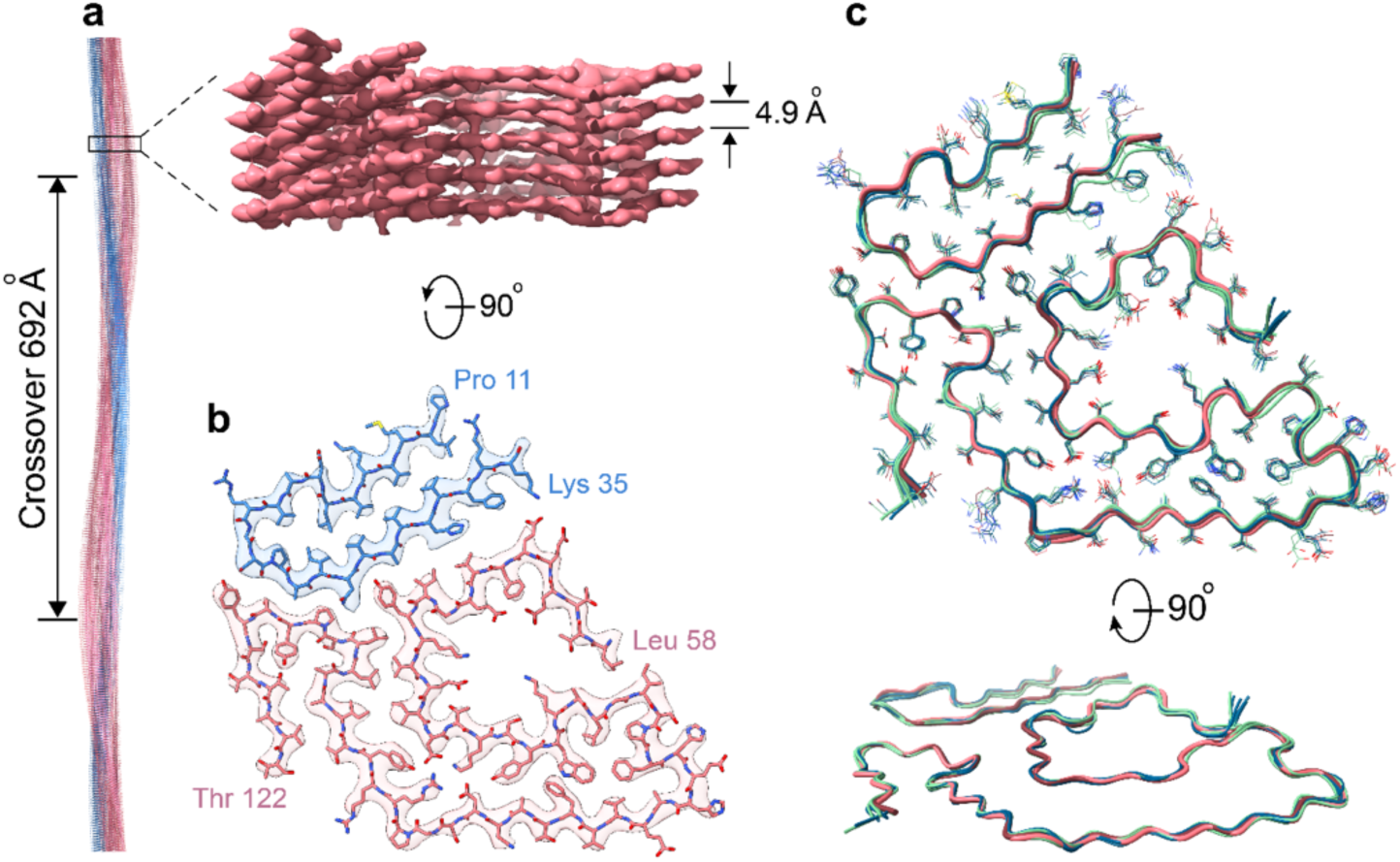
Cryo-EM structure of single fibril morphology from the heart of an ATTRv-V122Δ patient. **a.** Side view of the reconstructed fibril model showing the crossover distance, with the closeup side view of the map depicting the helical rise. **b.** Cryo-EM density and atomic model of the single fibril structure. The models contain two fragments of transthyretin colored blue (residues Pro 11 to Lys 35) and pink (residues Leu 58 to Thr 122). **c.** Structural backbone alignment of ATTRv-V122Δ fibrils (heart in pink and liver in dark brick) compared to ATTRwt (green) fibrils and a group of previously published ATTRv (blue) fibril structures, including ATTRv-V20I, ATTRv-V30M (heart and nerve), ATTRv-G47E, ATTRv-I84S, and ATTRv-V122I fibrils^16,17,23^. We have purposefully excluded polymorphic ATTRI84S fibrils from this alignment because our objective with this analysis is to show the similarity of this particular fibril morphology across multiple genetic backgrounds, with an r.m.s.d. value of 0.65 Å.

### CryoEM structure of N-to-C double fibrils

We determined the 3D structure of N-to-C double fibrils to a resolution of 3.7 Å (Supplementary Fig. 5). This structure comprises of two intertwined and staggered protofilaments. Optimal local symmetry search revealed a twist of -1.15° corresponding to a crossover distance of 764 Å (assuming a left-handed twist fibril) and the rise of 4.88 Å (Fig. 4a). Each of the two protofilaments adopts a conformation consistent with the single fibril model with minor modification due to a missing density at the N-terminal end. Thus, compared to the single fibril, the N-terminal fragment of the N-to-C double fibrils’ structure appears to be shorter, lacking the residues Pro 11 and Lys 35 (Fig. 4b). The interface of the two protofilaments is held by the hydrogen bonding interactions between Glu 92 to Arg 21’ and Asn 98 to Tyr 116’ (Fig 4e). To date, cardiac extracts of ATTR fibrils have been found to contain only single fibrils. In contrast, a recent study shows that the extract from the vitreous humor of an ATTRv-V30M patient can contain fibrils with single, double, and triple protofilaments^19^. In this study, the authors only determined the structure of the double fibril population; and within this subset, the inter-protofilament interactions involve the hydrogen bonding between His 90 to Glu 92’ and Glu 92 to His90’ (Fig. 4c, e). While the polar channel of N-to-C double fibrils’ structure adopts the same conformation as found in ATTRwt fibrils, variations were observed in the polar channel of ATTRv-V30M double fibrils from vitreous humor. In the latter, the residue Glu 62 flips 180° away from the solution and into the polar channel, and residues Glu 57 to Gly 67 (previously referred to as the polar channel gate) block the polar channel diagonally (Fig 4c, d).

**Figure 4:**
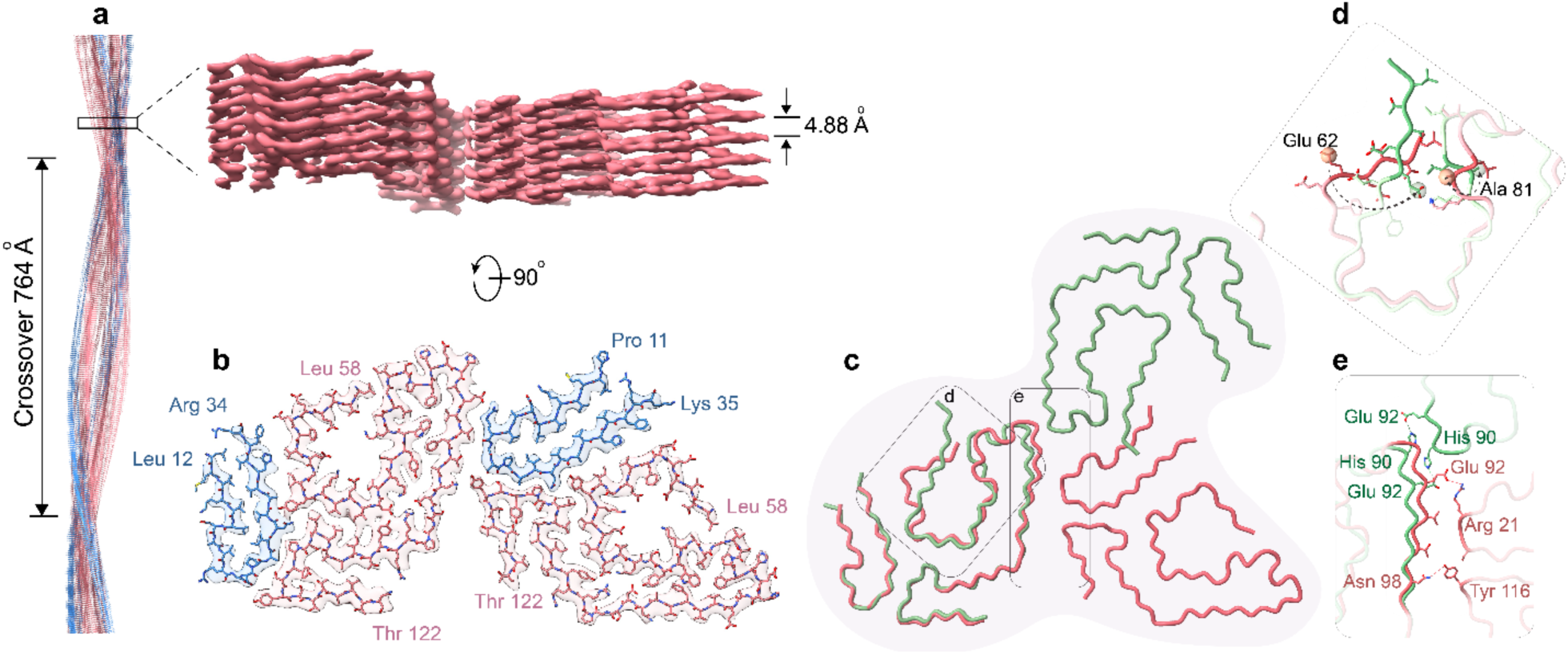
Cryo-EM structure of N-to-C double fibrils’ morphology from an ATTRv-V122Δ patient. **a.** Side view of the reconstructed fibril model of the N-to-C double fibril showing the crossover distance, with the closeup side view of the map depicting the helical rise. **b.** Cryo-EM density map and atomic model of the double fibril. On each protofilament, the model contains two fragments of transthyretin: the N-terminal fragment in blue (Leu 12 to Arg 34, and Pro 11 to Lys 35) and the C-terminal fragment in pink (Leu 58 to Thr 122). The two asymmetrically intertwined protofilaments’ interactions are mostly polar. **c.** Structural alignment of ATTRv-V122Δ (pink) and ATTRv-V30M eye (green) double fibrils highlighting the contrast of the symmetry of the double fibrils along with difference in residue interactions. **d**. Different arrangement of residues in the polar channel of the fibrils. **e**. Different interactions were observed at the fibrils’ interfaces of ATTRv-V122Δ (pink) and ATTRv-V30M eye (green).

### Cryo-EM structure of C-to-C double fibrils

The C-to-C double fibrils’ structure from the ATTRv-V122Δ amyloidosis patient was resolved at resolution of 4.3 Å. At this resolution, although we cannot see a discernable separation of the density map of individual layers within the fibril, we can identify in cross-sectional top view a two-fold symmetry (C2 symmetry) along the fibril axis. Because of the limited resolution of the density map, we chose not to construct an atomic model for the C-to-C double fibrils. Instead, we docked the model of the single fibril structure into this density map, with a C2 symmetry (Fig. 5b). We found that the predicted residues at the interface of the two fibrils engage in several interactions including two salt bridges between Glu 92 and Arg 104’, hydrophobic interactions between Val 94 and Pro 102’, and hydrogen bonds between Thr 96 and Ser 100’ (Fig. 5c).

**Figure 5:**
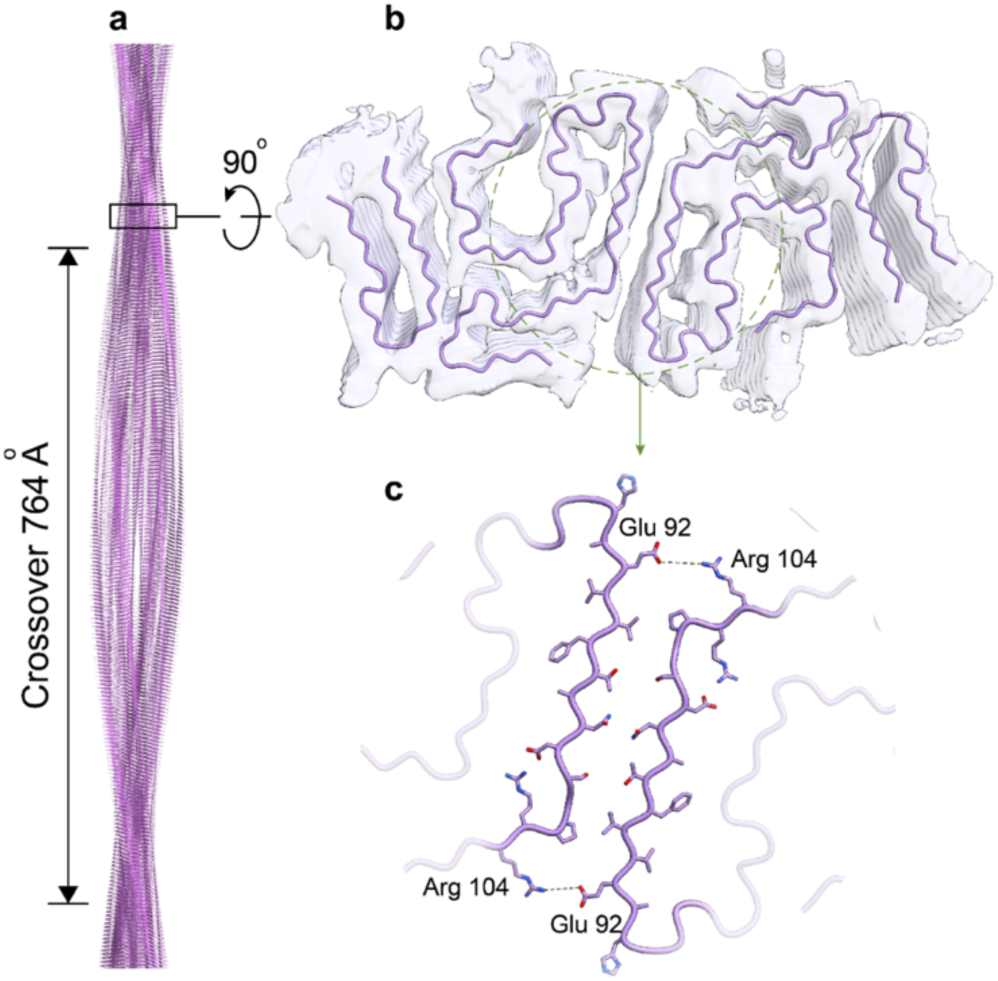
Cryo-EM structure of C-to-C double fibrils from an ATTRv-V122Δ patient. **a.** Cryo-EM model of the C-to-C double fibrils showing the crossover distance, with the closeup side view of the map depicting the helical rise. **b.** Docking illustrates the cryo-EM density model of the C-to-C double fibrils’ morphology showing the asymmetry of the fibrils. **c.** Docking indicating possibly polar interactions of fibrils are highlighted in purple.

### The two double fibrils’ structures can co-exist within the same fibril

We also traced the fibril segments that contributed to the different density maps of double fibrils’ morphologies present in the ATTRv-V122Δ cardiac extract, back to their corresponding micrographs. Our finding revealed a coexistence of segments from both N-to-C and C-to-C morphologies within the same fibril (Fig. 6a, b). Although, this phenomenon has been previously observed in published amyloid fibrils including those of systemic AL amyloidosis (subtype λ3 light chain)^25^ and ATTR amyloidosis (ATTRv-I84S)^18^, our study marks the first observation of this phenomenon within double fibrils.

**Figure 6:**
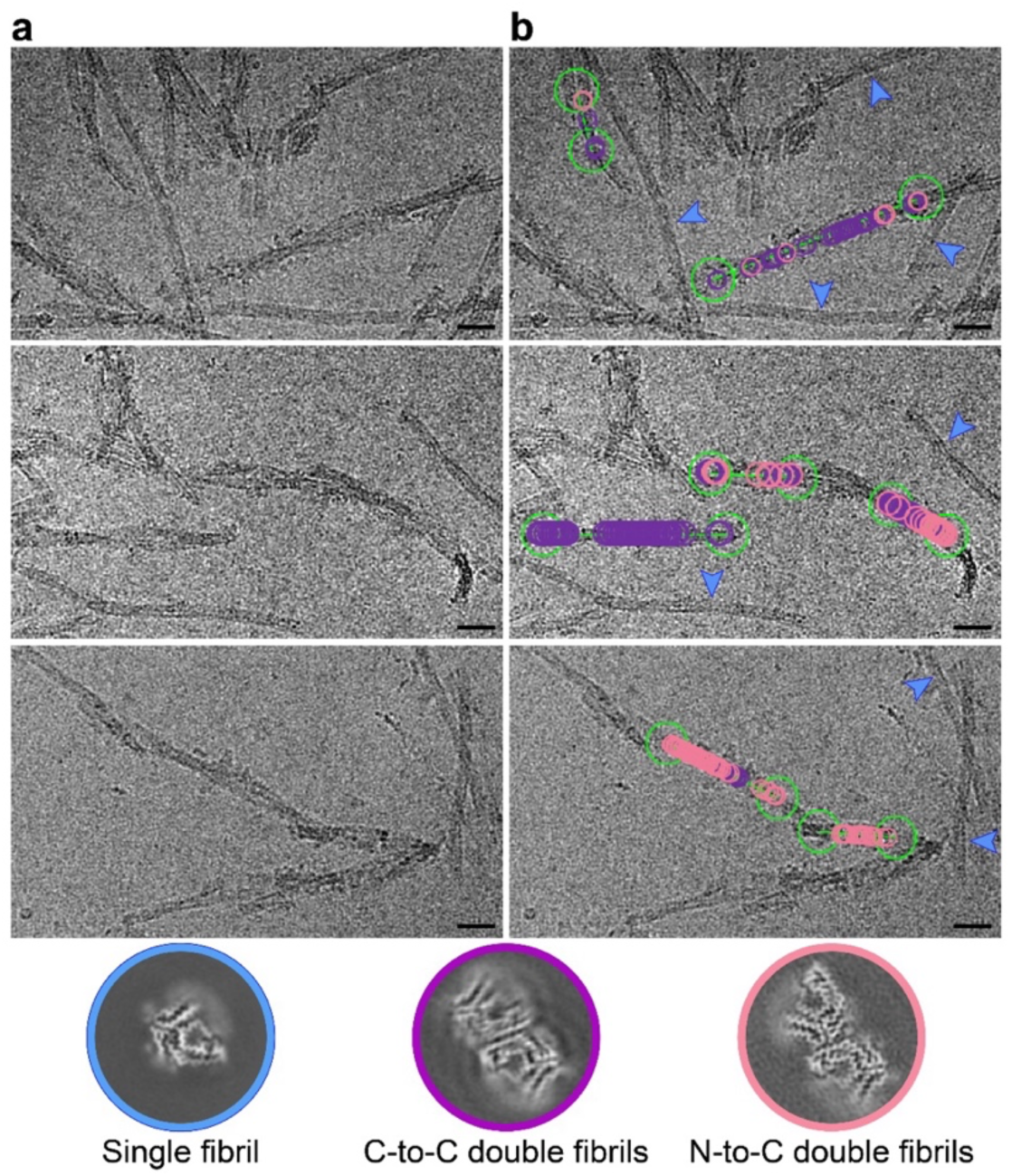
Existence of different morphologies within the same ATTRv-V122Δ fibril. **a.** Representative cryo-EM micrographs of cardiac ATTRv-V122Δ single and double fibrils. **b**. Representative cryo-EM micrograph tracing the single fibril (blue arrows), C-to-N double fibrils (purple circles), and C-to-C double fibrils (pink circles) back to the same fibril, indicating that the different morphologies can co-exist within the same fibril. Scale bar, 20 nm.

## Discussion

In this study, we determined the structure of *ex-vivo* ATTRv-V122Δ amyloid fibrils extracted from the heart and liver of a polyneuropathy patient. We found that these fibrils are polymorphic, as they are comprised of single and double protofilaments, in which each of the protofilaments adopts the same conformation similar to previously published ATTR fibrils including those from ATTRwt, ATTRv-V20I, ATTRv-P24S, ATTRv-A25S, ATTRv-V30M (from the heart and nerve), ATTRv-D38A, ATTRv-G47E, ATTRv-T60A (from the heart, kidney, thyroid, and liver), and ATTRv-I84S^16–18,23,24,26,27^. This broad structural similarity supports the hypothesis that a conserved fibril architecture may underlie more predictable clinical presentations such as ATTRwt cardiomyopathy. To date, only two mutations, ATTRv-I84S and ATTRv-V122Δ, exhibit structural polymorphism which may align with their predominantly polyneuropathic phenotypes. In conjunction with prior cryo-EM investigations on ATTR fibrils, our current findings open questions about a potential link between fibril structure and phenotype. Does fibril polymorphism play a role in the development of polyneuropathy in ATTR amyloidosis patients? If so, what mechanisms underlie the generation of this structural variability in neuropathy patients?

Early *in-vitro* studies offer the hypothesis that the phenotypic heterogeneity of ATTR amyloidosis may result from the differential effects of mutations in transthyretin aggregation. These studies describe how some pathogenic mutations are more prone to destabilizing the native protein conformation, facilitating tetramer dissociation into monomers and subsequent protein aggregation ^5,28^. Although there are no *in-vitro* studies on the stability of ATTRv-V122Δ tetramers to date, this variant may potentially be one example of a destabilizing mutation. Visualizing the crystal structure of tetrameric ATTRv-V122Δ (1bz8) in comparison to wild-type (4tlt), we indeed find an alternative conformation of the β-strand in where the mutation is located. Briefly, the absence of Val122 seems to induce a conformational alteration in subsequent residues at the C-terminal where the glutamines at position 127 reorient towards the transthyretin T4 binding pocket, potentially causing steric hindrance. This hindrance could prevent the binding of T4, destabilizing the unbound transthyretin tetramer and thereby promoting aggregation into amyloid^28,29^. It is conceivable to connect the differential effect of mutations in tetramer dissociation with the different onsets observed between genotypes. However, the association between this differential destabilization and the genotype-specific organotropism or the phenotypic variability observed within the same genotype is not so evident, and could be explained by other factors.

Cryo-EM studies of amyloid fibrils have yielded near-atomic resolution density maps revealing acute structural insights into amyloid disease pathologies and phenotypes for other conditions. Numerous structural examinations of amyloid fibrils isolated from the brains of patients afflicted with neurodegenerative diseases including tauopathies and synucleinopathies, consistently indicate a link between each phenotype and an amyloid structural fold^10–14^. On the contrary, ATTR amyloidosis displays a heterogeneous spectrum of fibril structures that are capable of co-existing within the same individual^18^. The most prominent structural differences are found in the number of protofilaments observed within the same fibrils and the conformation adopted by a polar channel that runs along the fibril axis. Regardless, different polymorphs can be found within the same fibril, indicating that they are capable of seeding each other (Fig. 6). The association of these fibril structures with phenotype in ATTR amyloidosis has not been established yet.

With the contribution of our study, we observe that, to date, primarily cardiomyopathic patients have one—and only one—fibril structure in cardiac tissue, whereas primarily polyneuropathic patients present a more variable structural landscape^16,18,19^. In ATTRwt patients and all primarily cardiomyopathic ATTRv patients, the polar channel appears in a closed state, with the residues 57 to 67 acting as a gate that locks the channel at a hydrophobic interface^16,17^. In contrast, in primarily polyneuropathic ATTRv-I84S patients, the gate that closes this channel suffers multiple perturbations, opening, exposing, or blocking the polar channel^18^. Additionally, in the eye of V30M patients, fibrils have multiple protofilaments and the gate is also perturbed^19^. Our study on ATTRv-V122Δ fibrils provide an additional polymorph where the gate remains closed, as shown in fibrils from cardiomyopathic patients, but the fibrils are made of single and double protofilaments, resembling those fibrils found in vitreous humor (Fig. 4c-e). Yet, the inter-protofilament interfaces found in cardiac ATTRv-V122Δ fibrils are not the same as the interface found in ATTRv-V30M fibrils from the vitreous humor (Fig. 4c-e).

The precise role of ATTR variants in shaping fibril polymorphism remains uncertain and requires further investigation. In the case of ATTRv-I84S, the mutation of residue 84 to serine could perturb a hydrophobic interface with the gate of the polar channel (around residue glycine 57), which could destabilize the “lock” of the gate that closes the polar channel, forcing and/or facilitating the formation of alternative gate conformations. However, the role of the V30M mutation is ambiguous as ATTRv-V30M fibrils from the vitreous humor also adopt an alternative gate conformation without seemingly destabilizing the gate lock^19^. Thus, the involvement of the V30M mutation in the formation of this alternative gate conformation or the formation of fibrils with multiple protofilaments in vitreous humor is not obvious. Similarly, the influence of the ATTRv-V122Δ deletion mutation in leading to multiple protofilaments remains unclear as the mutation site on the fibril is distant from any of the two potential inter-protofilament interfaces. Instead of *driving* the formation of new conformations, we surmise that the mutation may *allow for* the arrangement of the protofilaments into double fibrils. In other words, the shortening of one residue on the β-strand H of one protofilament could permit the interaction with another protofilament. This may apply to both double fibril morphologies reported from the ATTRv-V122Δ patient. Taken together, it is still unclear whether the V122Δ mutation impacts tetramer stabilization, drives the formation of particular fibril conformations, and/or affects protein aggregation and phenotype through alternative mechanisms.

Should there be a potential association between fibril polymorphism and polyneuropathy in ATTR amyloidosis, would this polymorphic variation be detectable, or imprinted, within the blood of ATTR patients? Recent studies from the Kelly lab and our lab indicate that transthyretin may aggregate in the blood of ATTR patients. The Kelly lab developed a peptide probe that detects non-native transthyretin species in the blood of neuropathic patients that are absent in those with cardiomyopathic or mixed phenotypes^30^. We, in turn, developed a peptide probe that detects big transthyretin aggregates in the blood of all ATTR patients, regardless of their phenotype^31^. These observations may implicate that the biology of ATTR amyloidosis with polyneuropathy may be different from that of ATTR amyloidosis with cardiomyopathy or mixed phenotypes. In neuropathic cases, the non-native species might serve as or evolve into polymorphic seeds, featuring single or multiple protofilaments. These seeds could then catalyze the polymerization of circulating wild-type and variant transthyretin into fibrils, also with single or multiple protofilaments, and/or adopting alternative gate conformations. Additional studies would be necessary in understanding how certain variants—and not others—contribute to the formation of polymorphic non-native transthyretin species or aggregates in the bloodstream.

The potential influence of chaperones and microenvironmental conditions on fibril structure should also be investigated. Although chaperones are traditionally known to play protective roles by facilitating protein folding and promoting clearance of protein misfolding, emerging evidence suggests that certain chaperones may generate seeding competent species^32,33^. If fibril seeds originate in the blood as speculated, chaperone interactions may play an important role in shaping their structures. Such interactions may modify seed architecture, which could influence fibril morphology as these seeds template fibril growth. In addition to chaperone effect, microenvironmental factors associated with differences in the medium where transthyretin circulates (blood versus vitreous humor, for instance) may further contribute to fibril polymorphism. For example, in vitreous humor of a patient with ATTRv-V30M amyloidosis, fibrils appeared with multiple protofilaments and with local variation at the polar channel, similar to those observed in fibrils from patients with ATTRv-I84S amyloidosis^18,19^. Understanding how specific variants, chaperone activity, and microenvironmental conditions drive the formation of polymorphic aggregates will be essential to understand clinical phenotypic variations in ATTR amyloidosis.

Given the evidence for structural polymorphism in ATTR fibrils, a key concern arises regarding how these structural variations might influence the efficacy and design of therapeutic treatments. Current treatments for ATTR amyloidosis do not directly target amyloid fibrils but rather focus on lowering transthyretin production or preventing tetramer dissociation. Of the five current FDA-approved drugs for ATTR amyloidosis, two of them are transthyretin tetramer stabilizers (tafamidis and acoramidis) and the other three are gene silencers (patisiran, vutrisiran, and inotersen)^34^. Since the action mechanism of these drugs target the upstream pathway prior to the fibril formation, amyloid structural polymorphism unlikely impact the efficacy of these treatments. However, fibril polymorphism could have significant implications for the development of future therapies that aim to disaggregate or remove amyloid deposits. For example, antibody-based “depleter” therapies currently in clinical trials^34^ may face challenges if polymorphism alters the accessibility or presentation of fibril epitopes. To address these challenges, future therapeutic strategies should focus on conserved structural regions within ATTR fibrils that remain consistent across different mutations and clinical phenotypes (cardiomyopathy, polyneuropathy, or mixed). These key conserved regions are the N-terminal segment spanning residues 11 to 35 and the C-terminal segment spanning residues 69 to 124. Targeting these conserved regions or shared structural motifs could minimize the effects of polymorphism and provide a unified therapeutic approach for all ATTR amyloidosis patients.

Our study has several limitations. First, due to the restricted availability of patient material in our biobank, we were only able to analyze heart and liver tissues from a single individual. Tissue from classical neuropathic sites such as sural nerve or skin was not available, preventing structural characterization of fibrils in regions directly related to polyneuropathy. Thus, the association between fibril polymorphism and neuropathic manifestations remains unresolved and warrants investigation using samples from additional patients. Second, ATTRv-V122Δ fibrils were isolated postmortem, meaning that the structures presented here likely represent end-stage fibrillar morphologies. Future studies incorporating longitudinal sampling or earlier disease-stage tissues will be essential to clarify how fibril structures evolve and how these polymorphisms relate to clinical heterogeneity in ATTR amyloidosis.

In summary, we structurally characterized cardiac and liver ATTRv-V122Δ fibrils from a polyneuropathy patient. We observed polymorphism within these fibrils, and we determined the structures of single fibril morphology in both the heart and liver and double fibril morphologies in the heart. Our study suggests that polymorphism in ATTR fibrils may manifest in predominantly polyneuropathic phenotypes. This study contributes to the understanding of the complex mutational landscape of ATTR amyloidosis and calls for further characterization of ATTR fibril polymorphism.

## Materials & Methods

### Patient description and tissue material

This investigation is based on the study of postmortem tissues from a 63-year-old Ecuadorian-origin man with the ATTRv-V122Δ mutation^20^. His medical history included a subarachnoid hemorrhage at age 57, followed by lower extremity weakness, numbness, and paresthesia. He also experienced significant peripheral and autonomic neuropathy, presenting as impotence, alternating constipation and diarrhea, urinary frequency, and difficulty walking. Amyloid deposits were identified in a rectal biopsy using Congo red staining and confirmed through immunohistological staining with an anti-TTR antibody. Echocardiography revealed hypertrophy of both the left and right ventricles, along with a refractile myocardial backscatter consistent with amyloidosis. The Office of the Human Research Protection Program exempted the study from Internal Review Board since the specimen was anonymized.

### Histologic confirmation of amyloidosis in human tissue

Fresh-frozen human tissue (heart and liver) was thawed at 4°C overnight, fixed in 20x volume excess of 10% neutral buffered formalin for 48 hours at room temperature with agitation, then transferred to 70% ethanol and paraffin processed. Sections were prepared for routine H&E staining, Congo Red, and Thioflavin-S. H&E staining was performed using Sakura DRS-601 robot and Leica-Surgipath Selectech reagents. Congo Red slides counterstained with hematoxylin were assessed for amyloid aggregation under bright-field. Thioflavin-S slides were evaluated for plaque and vascular amyloid under UV excitation (400-440 nm) and long-pass emission (470 nm+). Thioflavin-S staining involved sequential oxidation, bleaching, peroxidation, and acidification before incubating in alcoholic Thioflavin-S solution. Slides were then dehydrated, cleared, and cover slips affixed with Cytoseal 60 mounting media. This method is further described in the supplementary text.

### Fibril extraction from human cardiac and liver tissue

Amyloid fibrils were extracted from fresh-frozen human tissues, following a previously described protocol^16^. Briefly, ∼150 mg of frozen human tissue was thawed at room temperature, then finely cut into small pieces with a scalpel. The sample was washed with 1 mL Tris-calcium buffer (20 mM Tris, 150 mM NaCl, 2 mM CaCl_2_, 0.1% NaN_3_, pH 8.0), then centrifuged for 5 min at 3100 x g and 4 °C. The supernatant was removed, and the sample underwent four additional washes in Tris-calcium buffer. Following the washing steps, the pellet was resuspended in 1 mL of 5 mg/mL collagenase solution, then incubated overnight at 37 °C, shaking at 400 rpm. The next morning, the resuspension was centrifuged for 30 min at 3100 x g and 4 °C. The pellet was subsequently resuspended in 1 mL Tris-ethylenedianimenetetraacetic acid (EDTA) buffer (20 mM Tris, 140 mM NaCl, 10 mM EDTA, 0.1% NaN_3_, pH 8.0). This suspension was centrifuged for 5 min at 3100 x g and 4 °C, and this washing step with Tris-EDTA was repeated nine additional times. After these washing steps, the pellet was resuspended in 150 μL ice-cold water supplemented with 5-10 mM EDTA and centrifuged for 5 min at 3100 x g and 4 °C. This step was performed 3-5 additional times to release amyloid fibrils from the pellet, with EDTA aiding in fibril solubilization.

### Electron microscopy of negatively stained samples

Confirmation of amyloid fibril extraction was achieved through negative-stained transmission electron microscopy, following the described protocol^18^. In brief, a 3 μL sample was placed on a glow-discharged carbon film 300-mesh copper grid (Electron Microscopy Sciences), incubated for 2 min, and delicately blotted on filter paper to eliminate excess solution. Subsequently, the grid underwent negative staining with 5 μL of 2% uranyl acetate for 2 min, followed by gentle blotting to remove the staining solution. An additional 5 μL of uranyl acetate was applied and promptly removed from the grid. Specimens were examined using an FEI Tecnai 12 electron microscope with an accelerating voltage of 120 kV.

### Western blotting of extracted ATTR fibrils

Western blot analysis was conducted on the extracted fibrils to verify their type^35^. In summary, 0.5 µg of fibrils were dissolved in a tricine SDS sample buffer, boiled for 2 min at 85 °C, and then ran on a Novex™ 16% tris-tricine gel system using a Tricine SDS running buffer. The type of transthyretin was identified by transferring the gel contents onto a 0.2 µm nitrocellulose membrane and probing it with a primary antibody (1:1000) targeting the C-terminal region of the wild-type transthyretin sequence from GenScript. A secondary antibody, horseradish peroxidase-conjugated goat anti-rabbit IgG (dilution 1:1000, Invitrogen), was then used. Transthyretin content was visualized using Promega Chemiluminescent Substrate, following the manufacturer’s instructions.

### Amyloid seeding assay

Extracts of amyloid fibrils were used for seeding the formation of new fibrils from recombinant MTTR as detailed in our previous work^36^. In summary, the extracts were treated with 1% sodium dodecyl sulfate (SDS) and centrifuged for 1500 x g for 20 min in order to further purify the sample. This purification step was repeated twice, and the soluble fractions were removed. The sample then underwent three washes with sodium phosphate-EDTA (without the addition of 1% SDS) through centrifugation, and was then sonicated in a bath sonicator using cycled of 5 seconds on and 5 sec off for a duration of 10 min, using a minimum amplitude of 30 (Q700 sonicator, Qsonica). To measure the total protein content in the prepared seeds, the Micro BCA™ Protein Assay Kit (Thermo Fisher Scientific) was used in which a 2% (w/w) seed solution was added to 0.5 mg/mL recombinant MTTR in a final volume of 200 μL, containing 10 μM thioflavin T (ThT) and 1× PBS (pH 7.4). The ThT fluorescence emission was measured at 482 nm with absorption at 440 nm in a FLUOstar Omega (BMG LabTech) microplate reader. The plates (384 Well Optical Btw Plt Polybase Black w/o Lid Non-Treated PS, Thermo Fisher Scientific) were incubated at 37 °C with cycles of 9 min shaking (700 rpm double orbital) and 1 min rest throughout the incubation period. Measurements were taken every 10 min (bottom read) with a manual gain of 1000 fold. Confirmation of fibril formation was performed using transmission electron microscopy, as previously described.

### Mass Spectrometry (MS) sample preparation, data acquisition and analysis

0.5 µg of extracted ATTR fibrils were prepared for tryptic MS analysis by boiling in tricine SDS buffer and running on a Novex™ 16% tris-tricine gel. The gel was stained with Coomassie dye, and the ATTR smear was excised for MS analysis. Samples were digested overnight with trypsin, then cleaned up with solid-phase extraction before injection onto a Q Exactive HF mass spectrometer coupled to an Ultimate 3000 RSLC-Nano liquid chromatography system. MS scans were acquired at 120,000 resolution, with up to 20 MS/MS spectra obtained using higher-energy collisional dissociation. Data were analyzed using Proteome Discoverer v3.0 SP1, with peptide identification against the human reviewed protein database from UniProt. Raw data is deposited in the MassIVE data repository (accession number: MSV000094731). This method is further described in the supplementary text.

Raw MS data files were analyzed using Proteome Discoverer v3.0 SP1 (Thermo), with peptide identification performed using a semitryptic search with Sequest HT against the human reviewed protein database from UniProt. Fragment and precursor tolerances of 10 ppm and 0.02 Da were specified, and three missed cleavages were allowed. Carbamidomethylation of Cys was set as a fixed modification, with oxidation of Met set as a variable modification. The false-discovery rate (FDR) cutoff was 1% for all peptides. The mass spectrometry proteomics data have been deposited to MassIVE data repository, with accession number MSV000094731 ftp://MSV000094731@massive.ucsd.edu.

### Cryo-EM sample preparation, data collection, and processing

A 3.5 µL aliquot of the freshly extracted ATTR fibrils were applied to glow-discharged Quantifoil R 1.2/1.3, 300-mesh, Cu grids. The grid was then blotted with filter paper (100% humidity, 3 seconds blotting time and -1 blot force) to remove the excess sample, and plunge frozen into liquid ethane using a Vitrobot Mark IV (FEI). This Cryo-EM sample was screened on the Talos Arctica at the Cryo-Electron Microsopy Facility (CEMF) at University of Texas Southwestern Medical Center (UTSW). Cryo-EM data was collected on a 300 kV Titan Krios microscope (FEI) at the Stanford-SLAC Cryo-EM Center (S^2^C^2^) (Supplementary Table 1). All movies were recorded using a Falcon4 camera and EPU software (Thermo Fisher Scientific). All collection parameters are detailed in Supplementary Table 1. Raw movies were gain-corrected, aligned and dose-weighted using RELION’s own implemented motion correction program^37^. Contrast transfer function (CTF) estimation was performed using CTFFIND 4.1^38^. All steps of helical reconstruction, three-dimensional (3D) refinement, and post-process were carried out using RELION 4.0^22^. Single filaments were picked automatically using Topaz, while double filaments were picked manually ^39^. Particles were initially extracted using a box size of 768 pixels, then down-scaled to 256 pixels (with an inter-box distance of ∼10% of the box length) to determine the cross-over distance and the population distribution for each morphology. To further process the data, single filaments and double filaments were extracted in box of 192 pixels and 320 pixels, respectively. Reference-free 2D classifications were performed for all datasets for three rounds with 25 iterations each round to select suitable particles for 3D reconstructions. Fibril helices were assumed to be left-handed.

### Reconstruction of single filaments

We used an elongated Gaussian blob as an initial reference for 3D classifications of single filaments for both heart and liver samples. The best looking class from these 3D classifications was used as a reference (low-passed to 10Å) for Refine3D jobs. A full mask was used in subsequent rounds of 3D classification without image alignment where particles were split into 3 classes (or ∼ 30,000 particles/class). Subsequent rounds of the 3D reconstruction were performed using Refine3D jobs with a full mask. Bayesian particle polishing and multiple rounds of CTF refinements were carried out until no further improvement in map resolution was observed. Final maps were post-processed using the recommended standard procedures in RELION, and the overall resolutions were estimated at threshold of 0.143 in the Fourier shell correlation (FSC) curve between two independently refined half-maps (Supplementary Fig. 4).

### Reconstruction of double filaments

Only the cardiac double filaments were reconstructed, because the liver sample did not have sufficient number of particles for reliable reconstruction. For 3D classifications of cardiac double filaments, the initial model was generated from 2D class average images using a RELION built-in, script relion_helix_inimodel2d^40^. We performed 3D classifications to select the best particles leading to the best reconstructed maps. Subsequent 3D auto-refinements and CTF refinements were carried out to obtain higher resolution maps. Final maps were post-processed using the recommended standard procedures in RELION, and the overall resolutions were estimated at threshold of 0.143 in the Fourier shell correlation (FSC) curve between two independently refined half-maps (Supplementary Fig. 4).

### Stabilization energy calculation

The stabilization energy per residue was calculated by the sum of the products of the area buried for each atom and the corresponding atomic solvation parameters^41^ (Supplementary Fig. 6). The overall energy was calculated by the sum of energies of all residues, and assorted colors were assigned to each residue in the solvation energy map.

### Model building

The refined maps of the different morphologies found in this patient were post-processed in RELION before building their models^42^. We used our previously published model of ATTRI84S (PDB code 8tdn) as the template to build the model of ATTRv-V122Δ single fibril. This model was then used as a template for building the double fibrils’ models. Residue modification, rigid body fit zone, and real space refine zone were performed to obtain the resulting models using COOT v0.9.8.1^43^. All the statistics are summarized in Supplementary Table 1.

### Filament tracing

The fibril segments’ coordinates (rlnCoordinateX and rlnCoordinateY) from the final Refine3D were extracted from the relion file run_data.star. These coordinates were used to create XY-graphs, which were then adjusted to align with the dimensions of the respective micrographs. Of note, only the clearly defined fibril segments were considered for reconstruction, excluding any subpar segments. From this method, the positions of these clearly defined segments, particularly when they belong to the same fibril, may appear as intermixed as depicted in Fig. 6.

## Acknowledgments

In recognition of the contributions made by the Late Dr. Merril D. Benson, who significantly advanced the understanding of amyloid diseases and provided support to affected families for decades. Special thanks to the patient and family who generously donated tissues, as well as the University of Indiana for providing the material. We thank the UTSW Cryo-Electron Microscopy Facility, the UTSW Structural Biology Laboratory, the UTSW Electron Microscopy Core Facility, the national cryo-EM facilities Stanford-SLAC (project CA60) for instrumentation, technical support, and/or data collection. We thank the UTSW Proteomics core for technical assistance with the proteomics experiments.

## Contributions

Conceptualization: L.S., Y.A., B.A.N. Methodology: L.S., Y.A., B.A.N, C.K., V.S., S.A. and Investigation: Y.A., B.A.N., C.K., V.S., S.A., J.M.S., C.L.E., B. M. E., R.P., M.P., M.C.F.R., P.B., P.S., and L.W. Visualization: B.A.N., L.S., and Y.A. Funding acquisition: L.S. Project Administration: L.S. Supervision: L.S. Writing—original draft: Y.A., L.S., and B.A.N. Writing—review and editing: L.S, Y.A, B.A.N., and P.B.

## Funding

American Heart Association, Career Development Award 847236, L.S.

National Institutes of Health, National Heart, Lung, and Blood Institute, New Innovator Award DP2-HL163810, L.S.

Welch Foundation, Research Award I-2121-20220331, L.S.

UTSW Endowment, Distinguished Researcher Award from President’s Research Council and start-up funds, L.S.

Junta de Andalucía, EMERGIA20_00276, R.G.P.

Cryo-EM research was partially supported by the following grants:

National Institutes of Health grant U24GM129547, Department of Energy Office of Science User Facility sponsored by the Office of Biological and Environmental Research

Department of Energy, Laboratory Directed Research and Development program at SLAC National Accelerator Laboratory, under contract DE-AC02-76SF00515

NIH Common Fund Transformative High Resolution Cryo-Electron Microscopy program (U24 GM129539)

The Cryo-Electron Microscopy Facility and the Structural Biology Laboratory at UTSW are supported by a grant from the Cancer Prevention & Research Institute of Texas (RP170644).

The Electron Microscopy Core Facility at UTSW is supported by the National Institutes of Health (NIH) (1S10OD021685-01A1 and 1S10OD020103-01).

Part of the computational resources were provided by the BioHPC supercomputing facility located in the Lyda Hill Department of Bioinformatics at UTSW. URL: https://portal.biohpc.swmed.edu.

