## Supplementary Figures and Tables for "Amyloid fibril polymorphism in the heart and liver of an ATTR amyloidosis patient with polyneuropathy attributed to the ATTRv-V122Δ variant"

**Affiliations:**
*[1]Center for Alzheimer's and Neurodegenerative Diseases, Department of Biophysics, Peter O'Donnell Jr Brain Institute, University of Texas Southwestern Medical Center (UTSW), Dallas, TX, USA.*
*[2]Department of Pathology, University of Texas Southwestern Medical Center (UTSW), Dallas, TX, USA.*
*[3]Department of Ophthalmology, University of Texas Southwestern Medical Center (UTSW), Dallas, TX, USA.*
*[4]Department of Pathology and Laboratory Medicine, Indiana University School of Medicine, Indianapolis, IN, USA.*

**Supplementary Figures:**

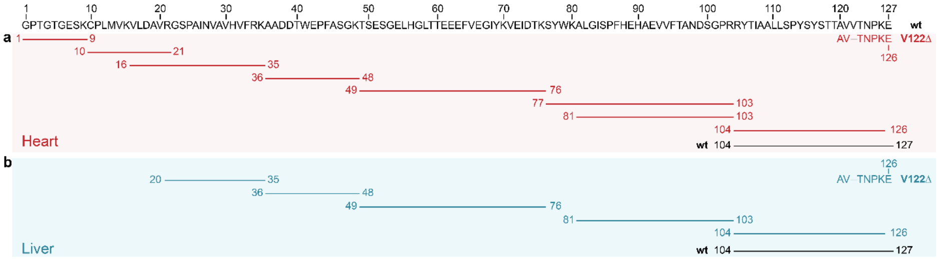


**Supplementary Figure 1:** Findings obtained from semi-tryptic mass spectrometry analysis of the heart and liver ATTRv-V122Δ fibril samples. Both the full length wild-type and mutant transthyretin were detected in these samples. The results confirm the presence of the V122Δ mutation.

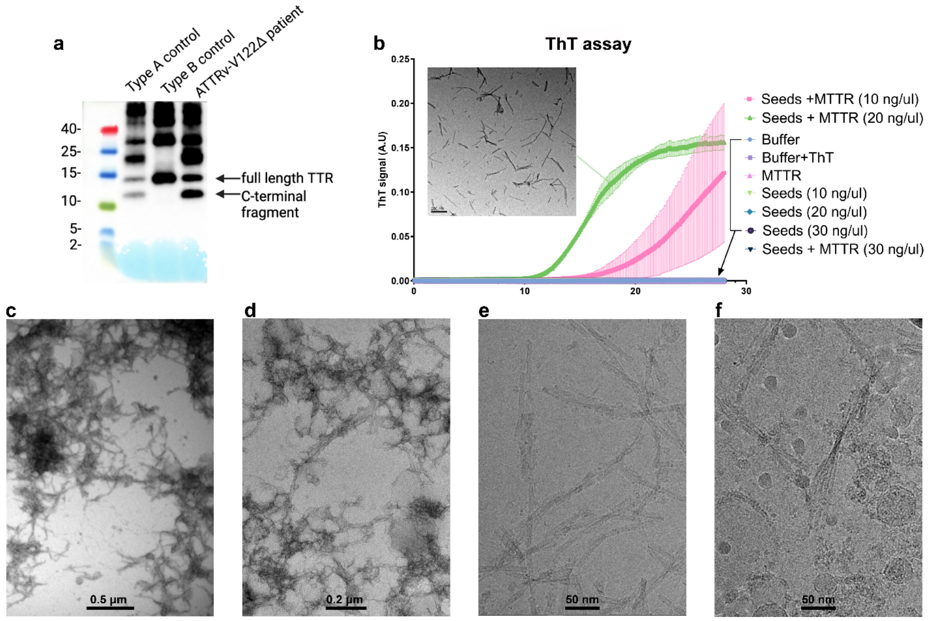


**Supplementary Figure 2: Confirmation of fibril extraction from the heart and liver of an ATTRv-V122Δ patient. a** Anti-TTR western blot of extracted cardiac fibrils (0.5 µg) using an antibody that identifies transthyretin fragmentation of the C-terminus, confirming the Type A nature of these fibrils. **b** Assessment of seeding activity through Thioflavin T assay demonstrating the seeding competence of ATTRv-V122Δ fibrils, with a negative stained image of seeded fibrils. **c and d** Representative negative stained images of extracted fibrils from the heart and the liver showing an abundance of fibrils, respectively. Scale bar, 0.5 µm and 0.2 µm, respectively. **d** Representative cryo-EM micrograph of fibrils from the heart and the liver of the ATTRv-V122Δ patient. Scale bar, 50 nm.

**Commented [AN1]:** Need cryoEM from Liver dataset

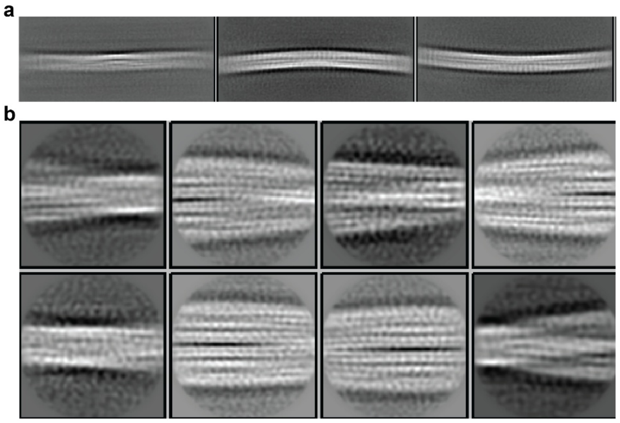


**Supplementary Figure 3: from the ATTRv-V122Δ patient. a and b are** representative 2D class averages of single fibril and double fibrils, respectively. Single and double fibrils were at particle box size of 1024 pixel and 300 pixels, respectively.

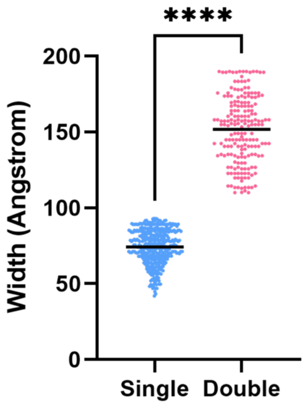


**Supplementary Figure 4.** Width measurements of both the single filament and double filaments' morphologies.

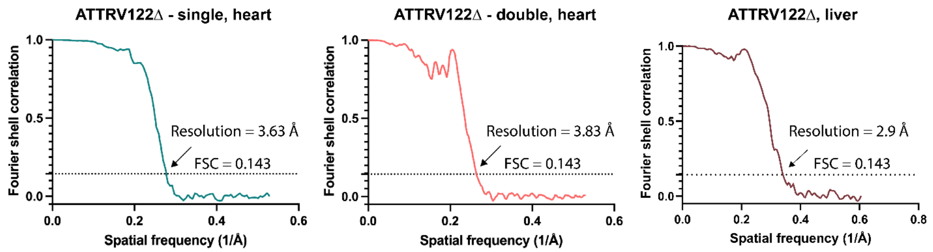


**Supplementary Figure 5. FSC curves.** Assessing the resolution of cryo-EM maps through Fourier shell correlation (FSC) curves from two separately refined half-maps of ATTRv-V122Δ fibrils. FSC curves are shown for the heart single filament and double filament, and the liver single filament.

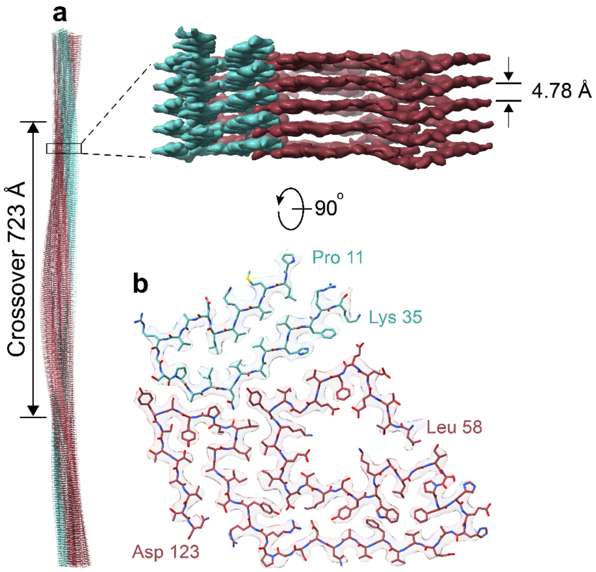


**Supplementary Figure 6. Cryo-EM structure of single fibril morphology from the liver of an ATTRv-V122Δ patient. a.** Side view of the reconstructed fibril model showing the crossover distance, with the closeup side view of the map depicting the helical rise. **b.** Cryo-EM density and atomic model of the single fibril structure. The models contain two fragments of transthyretin colored blue (residues Pro 11 to Lys 35) and pink (residues Leu 58 to Asp 123).

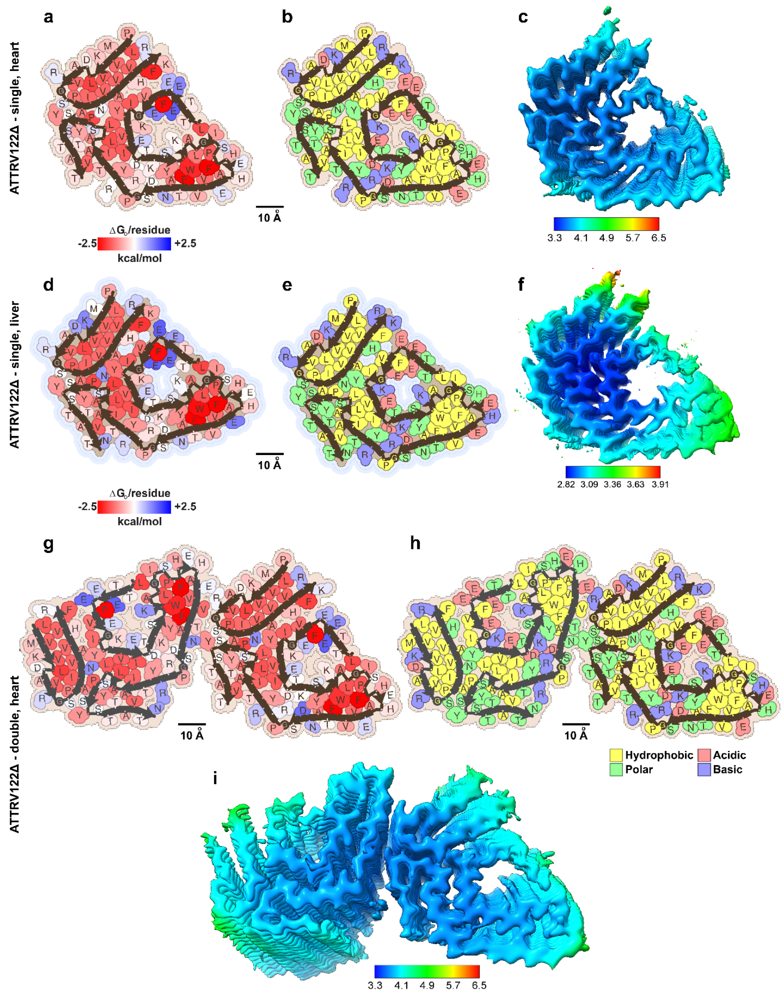


**Supplementary Figure 7. Structural analysis of ATTRv-V122Δ fibril. (a-c)** Solvation energies per residue, schematic residue composition and the local resolution map for the ATTRv-V122Δ – single filament from the heart, respectively. **(d-f)** Corresponding analyses for the ATTRv-V122Δ – single filament from the liver. **(g-i)** Corresponding analyses for the ATTRv-V122Δ – double filament from the heart. In the solvation energy panel, residues are color-coded from favorable (red, -2.5 kcal/mol) to unfavorable (blue, 2.5 kcal/mol) stabilization energy. In the residue composition panels, residues are color-coded by their amino acid category, as indicated by the labels. Scale, 10 Å.

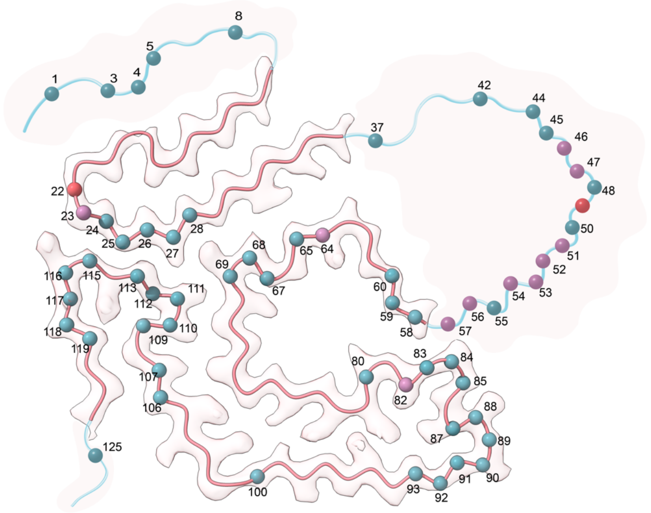


**Supplementary Figure 8.** Comparison of possible proteolytic sites on ATTRv-V122Δ and ATTRwt fibrils based on semi-tryptic peptides from MS data. The colored balls represent proteolytic sites on ATTRv-V122Δ fibrils only (salmon pink), on ATTRwt fibrils only (blue) and on both fibrils (purple).

**Supplementary Table 1.** Data collection and model refinement statistics.

| | Heart | | | Liver |
|---|---|---|---|---|
| **Data collection of ATTRv-V122Δ** | **Single filament** | **N-to-C Double filament** | **C-to-C Double filament** | **Single filament** |
| **Microscope (Model)** | Titan Krios (G3i) | Titan Krios (G3i) | Titan Krios (G3i) | Titan Krios (G2) |
| **Acceleration Voltage (kV)** | 300 | 300 | 300 | 300 |
| **Detector** | Falcon 4i | Falcon 4i | Falcon 4i | Falcon 4i |
| **Software** | EPU 3.5 | EPU 3.5 | EPU 3.5 | SerialEM 4.1.5 |
| **Magnification** | 130,000x | 130,000x | 130,000x | 105,000x |
| **Pixel size at detector (Å/px)** | 0.946 | 0.946 | 0.946 | 0.413 |
| **Defocus range (µm)** | 0.9 to 2.1 | 0.9 to 2.1 | 0.9 to 2.1 | 1 to 2.4 |
| **Total dose (e)** | 40 | 40 | 40 | 50 |
| **Exposure time (sec)** | 3.67 | 3.67 | 3.67 | 2.06 |
| **Number of movie frames** | 40 | 40 | 40 | 50 |
| **Usable micrographs** | 5759 | 5759 | 5759 | 4366 |
| **Number of segments after 2D**<br>Box size = 768 pixel | 899722 | 152423 | | n/a |
| **Total extracted segments**<br>Box size (pixel) | 498436<br>(192) | 726223<br>(320) | 726223<br>(320) | 345559<br>(300) |
| **Number of segments after 2D**<br>Box size (pixel) | 355380<br>(192) | 430731<br>(320) | 430731<br>(320) | 104042<br>(300) |
| **Number of straight segments** | 21641 | 0 | 0 | 7580 |
| **Number of segments after 3D**<br>Box size (pixel) | 32315<br>(192) | 40635<br>(320) | 120296<br>(256) | 21384<br>(300) |
| **Symmetry imposed** | C1 | C1 | C1 | C1 |
| **Helical rise (Å)** | 4.9 | 4.88 | 4.88 | 4.82 |
| **Helical twist (°)** | -1.27 | -1.15 | -1.15 | -1.19 |
| **Crossover length (Å)** | 692 | 764 | 764 | 729 |
| **B factor** | -117.3 | -97.2 | -189 | -49.97 |
| **Map resolution (Å; FSC=0.143)** | 3.63 | 3.82 | 4.3 | 2.95 |
| **Map resolution (Å; FSC=0.5)** | 4.3 | 4.4 | 4.7 | 3.4 |
| **Non-hydrogen atoms** | 3555 | 7030 | n/a | 3575 |
| **Protein residues** | 455 | 895 | n/a | 455 |
| **Number of chains** | 5 | 10 | n/a | 5 |
| **Water/ligands** | 0/0 | 0/0 | n/a | 0/0 |
| **MolProbity score** | 1.84 | 1.73 | n/a | 1.86 |
| **Clash score** | 9.86 | 8.21 | n/a | 13.45 |
| **Rotamer outliers (%)** | 0.00 | 0.00 | n/a | 0.00 |
| **R.M.S deviations bonds (Å)** | 0.003 | 0.003 | n/a | 0.003 |
| **R.M.S deviations angle (°)** | 0.537 | 0.644 | n/a | 0.667 |
| **Ramachandran plot** (%)<br>Favored<br>Allowed<br>Outliers | <br>95.40<br>4.60<br>0.00 | <br>95.91<br>4.09<br>0.00 | n/a | <br>96.55<br>3.45<br>0.00 |
| **CaBLAM outliers (%)** | 3.61 | 4.91 | n/a | 4.82 |
| **Model vs Data** | 0.79 | 0.81 | n/a | 0.84 |
